# SARS-CoV-2 has not emerged in roe, red or fallow deer in Germany or Austria during the COVID 19 pandemic

**DOI:** 10.1101/2022.02.18.480872

**Authors:** Andres Moreira-Soto, Christian Walzer, Gábor Á. Czirják, Martin Richter, Stephen F. Marino, Annika Posautz, Pau Rodo de Yebra, Gayle K. McEwen, Jan Felix Drexler, Alex D. Greenwood

**Author notes:** These authors were co–principal investigators (Jan Felix Drexler, Alex D. Greenwood). ^*^**Correspondence**: Alex D. Greenwood, Department of Wildlife Diseases, Leibniz Institute for Zoo and Wildlife Research, Alfred-Kowalke-Str.17, 10315, Berlin, Germany.

## Abstract

Spillover of SARS-CoV-2 to North American white tailed deer (*Odocoileus virginianus)* has been documented. We evaluated pre and pandemic exposure of German and Austrian deer species using a SARS-CoV-2 pseudoneutralization assay. In stark contrast to North American white tailed deer, we found no evidence of SARS-CoV-2 exposure.

**Article Summary Line:** Using a sensitive serological assay, 433 pre and pandemic deer samples from Germany and Austria tested negative for SARS-CoV-2-specific antibodies highlighting a major difference between central European and North American deer exposure and in their epidemiologic roles.

## The Study

The COVID-19 pandemic caused by the SARS-CoV-2 virus has spread from humans to domestic animals, captive and free ranging wildlife (1–5). In some cases, the result was the spillback of a novel SARS-CoV-2 variant to humans, making new wildlife reservoirs a clear threat to human health. Spillover is of particular concern in North America where there is strong evidence that white tailed deer (*Odocoileus virginianus*) have been infected in the U.S. and Canada with prevalences reaching 82.5% (3). White tailed deer may now represent a reservoir in which SARS-CoV-2 may evolve and potentially spill back to humans with unpredictable health consequences. White tailed deer are social animals that often form large herds, are important game species and often live in peri-urban and urban environments, making them unsurprising candidates as a viral reservoir species for pathogens with zoonotic potential. However, it is unclear how generalized spillback is in deer. In Europe, multiple species of deer are heavily managed and hunted but hunting legislation and management practices differ substantially between the U.S. and Europe, within Europe and even within individual countries and regions. Different deer species may also vary in the sequence of their ACE2 receptor (the receptor with which the SARS-CoV-2 virus spike protein interacts), which could hinder or promote infection (6).

To assess whether SARS-CoV-2 spillover has occurred in Europe, we applied SARS-CoV-2 pseudoneutralization assays to pandemic collected sera from the three main species of European cervids, namely red deer (*Cervus elaphus*) (n=67), roe deer (*Capreolus capreolus*) (n=94) and fallow deer (*Dama dama*) (n=68) (Table 1). Samples were collected from Germany and Austria, two countries with long deer management traditions. Sample size calculations required a minimum of 81-138 animals, assuming seroprevalences ranging from 30% to 90%, a large population size and 95 % confidence intervals (calculations were performed using the package epiDisplay in R). To assess assay specificity, we further tested 204 pre-pandemic sera. In stark contrast to North American white tailed deer, no pre or pandemic collected deer tested were positive for SARS-CoV-2 antibodies (Table 1, Supplementary Table). All positive and negative controls were in the 450 nm range stated in the manufacturer’s instructions. The assay cutoff for the cPass SARS-CoV-2 Neutralization Antibody Detection Kit is 30% inhibition (Supplmental Figure 1).

**TABLE 1.** Number and origin of samples from each deer species analyzed for anti-SARS-CoV-2 antibodies

|  | Germany |  | Austria |
| --- | --- | --- | --- |
|  | pre pandemic years | pandemic years | pandemic years |
| Roe deer | 74 | 94 |  |
| Red deer | 45 | 16 | 51 |
| Fallow deer | 85 | 68 |  |
| Total | 204 | 178 | 51 |

\*A SARS-CoV-2 surrogate virus neutralization test (sVNT; GenScript, USA), which measures antibody-mediated inhibition of SARS-CoV-2 receptor binding domain (RBD)-ACE2 interaction was used to test pre- and pandemic samples (6). Briefly, equal volume of deer sera were pre-incubated with HRP-conjugated RBD for 30 min at 37 °C, followed by addition to ACE2-coated ELISA plate for 15 min at room temperature. Colorimetric signal was developed using TMB substrate and stopped with 1 M HCl. Absorbance readings at 450 nm were acquired using a microplate reader. This test achieves 99.93% specificity and 95–100% sensitivity and the species-independent performance has been validated in rabbit, ferret, mice, cats, dogs, hamsters and alpaca serum (7, 8). The same assay was employed in North American white tailed deer with high seroprevalence (2).

**Figure 1.**
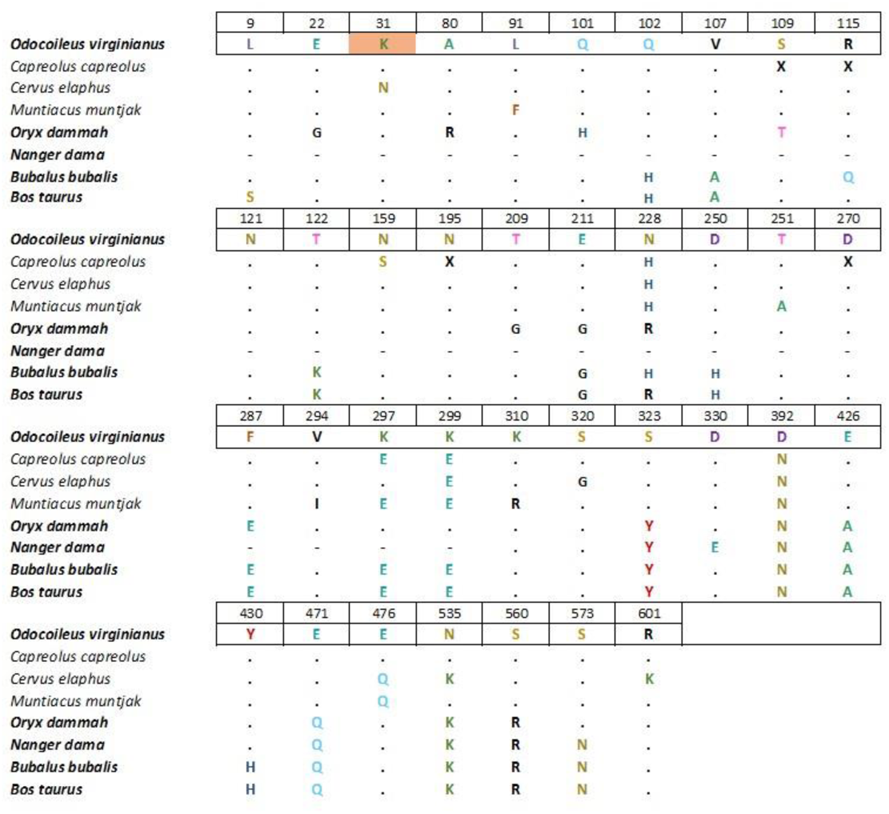
Multiple sequences the ACE2 protein were performed with the species characterized in (5) and additional deer species included from GenBank (GCA_000751575.1, XP_037678579.1, XP_012949915.3, QEQ50331.1, KAB0345583.1, XP_043752042.1, XP_036696353.1). The full alignment is in Supplemental Figure 1. The figure only shows deer and related bovids and only positions where there are amino acid changes relative to white tailed deer (*Odocoileus virginianus*). Alignments were performed using ClustalW Multiple Alignment in BioEdit version 7.2.5. The names of species highlighted in bold correspond to species known to be susceptible to SARS-CoV-2 and the amino acid position highlighted in orange is a hot spot for SARS-CoV-2 S-binding (5). Ambiguities in database sequences are designated as X. Dots represent identity to the reference and dashed represent missing sequence information.

We also compared the ACE2 receptor sequences for all deer and related bovids in the databases including roe and red deer to determine if they have amino acid changes that would be consistent with resistance to infection with SARS-CoV-2. Alignment of the ACE2 receptor among deer and related bovids showed that with the exception of K-N substation in the red deer relative to white tailed deer, none of the sequenced deer species had any substitutions that distinguished them from SARS-CoV-2 susceptible cervid species (Figure, Supplementary Figure 2).

The contrast between our results and the multiple reports of high prevalence infection in North American white tailed deer may have several explanations. First, a single relevant amino acid change was observed in red deer at position 31 (K-N) in the ACE2 receptor compared to other deer and bovids. While this could in principle explain infection resistance in red deer, roe deer were identical at all sites of import to white tailed deer yet were not infected in our study. It is not clear if this difference would make red deer resistant as other susceptible species vary at this position (6, Supplemental Figure 2).

In Europe the historic feudal systems, land use, and land allocation practices shaped hunting traditions. Across Germany, Austria and a large part of Switzerland hunting is implemented using a district-based hunting system (Revier) and wildlife belongs to no-one (*res nullius*) (9). In contrast, in North America wildlife is considered a public resource managed by the government independent of the land or water where wildlife lives and tightly linked to wildlife conservation (*res communis*). This distinction is important in terms of the extent to which government bodies can control and influence the management of wildlife and hunting practices. Therefore, in Europe deer management is more local than in North America (10).

Across the USA and Canada, the deer hunting season is generally shorter, lasting 3-4 months while in Europe it varies widely, from less than four weeks in some areas of Switzerland to 213-300 plus days in Austria and Germany (11). Hunting practices vary across the two regions: in North America deer hunting over bait and attractants is allowed in some 20 plus states, while in many regions of Europe this practice is either prohibited or frowned upon (1). Across North America feeding practices range widely by region from allowed, to strongly discouraged, to illegal. In continental Europe, supplementary seasonal feeding of deer is a common practice associated with, maintaining, in part, high densities and decreasing damage to agriculture and forestry assets (9, 12). However, the “Revier” structure maintains these concentrations locally and in rural areas. A major difference between white - tailed deer and red deer with respect to SARS-CoV-2 spillback from humans is that the former thrives seasonally at high densities in urban and peri-urban landscapes (13).

## Conclusions

A combination of extensive versus limited feeding, which could expose animals to human exudates in contaminated feed, and high density in urban and peri-urban landscapes that could expose herds to human waste, may be the key difference in SARS-CoV-2 spillback to deer populations. The lack of urban or peri urban populations and the (Revier) management system likely limits, thus far, contacts among central European deer populations may diminish human-wildlife interactions sufficiently to prevent the spread of infection. In this context, it would be advisable in Europe to maintain and strengthen barriers to spread, for example, to ensure that deer kept in wildlife parks have no contact with free ranging deer and prevent deer from settling near towns and villages to minimize future spillovers. In North America, it appears important to address behaviours related to feeding of deer in urban settings and revisiting feeding practices in deer management and hunting. Maintaining natural barriers will help prevent future spillovers and spillback of SARS-CoV-2 between deer and human populations (14). The consequences of inaction could be further spillback to humans of viral variants evolving in novel hosts as has occurred in Danish and Dutch mink farms during the pandemic (4,5).

## Supporting information

Sample metadata for samples analyzed in the study

Summary of serological results

Full alignment of cervid and other mammalian ACE2 receptor sequences

## Acknowledgements

The authors thank the help of hunters and colleagues involved in the collection of the archived and recent samples. The authors wish to thank Karin Hönig and Katja Pohle for their laboratory support.

## Biographical Sketch (first author)

Andres Moreira-Soto, DVM, PhD. is a senior postdoc in Charité University hospital, Berlin. His research focuses on newly emerging viral diseases in humans and animals.

## Notes

### Competing Interest Statement

The authors have declared no competing interest.

