## Supplementary figures and images for "SARS-CoV-2 has not emerged in roe, red or fallow deer in Germany or Austria during the COVID 19 pandemic"

### Summary of serological results

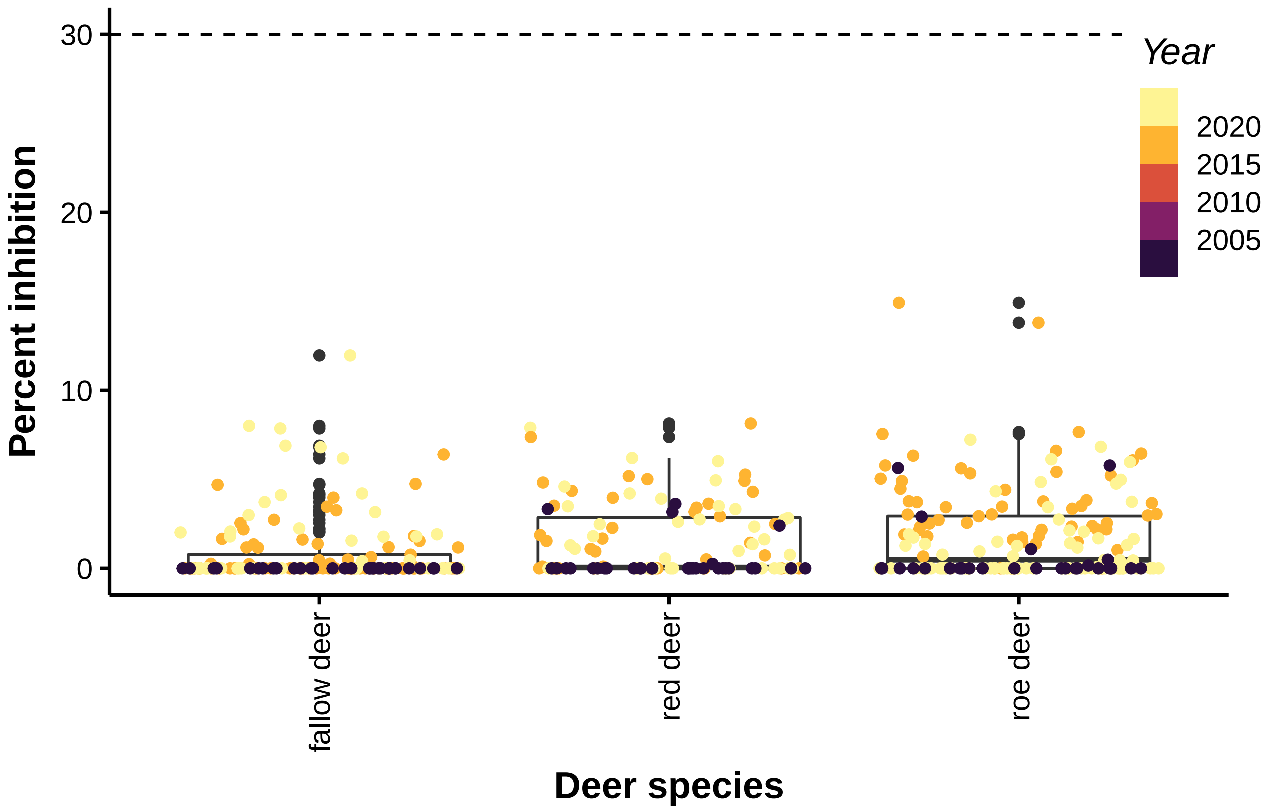
