## Supplementary material for "SARS-CoV-2 has not emerged in roe, red or fallow deer in Germany or Austria during the COVID 19 pandemic": Full alignment of cervid and other mammalian ACE2 receptor sequences

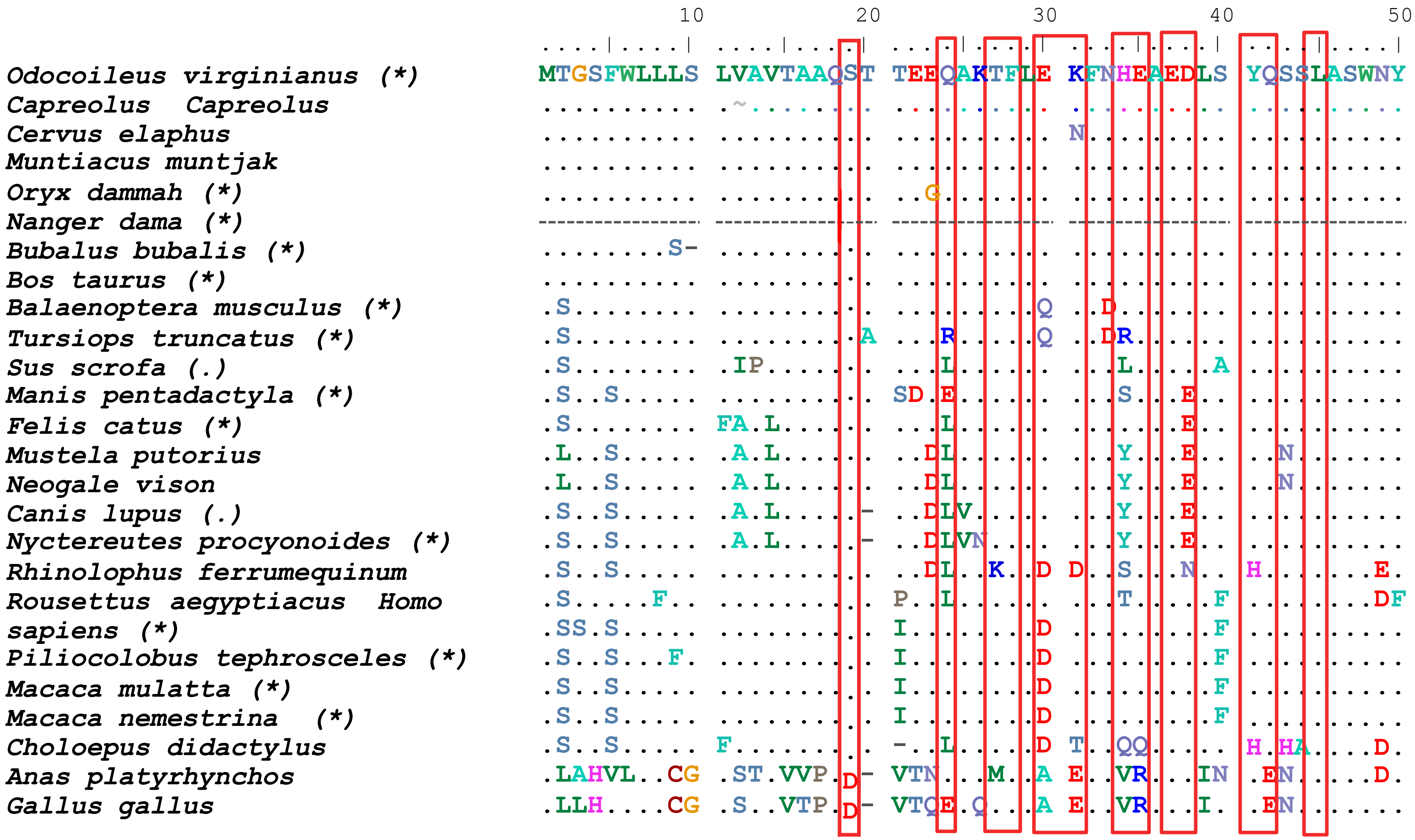
10
20
30
40
50
Odocoileus virginianus (*)
MTGSFWLLLS LVAVTAAQST TEEQAKTFLE KFNHEAEDLS YQSSLASWNY
Capreolus Capreolus
Cervus elaphus
Muntiacus muntjak
Oryx dammah (*)
Nanger dama (*)
Bubalus bubalis (*)
Bos taurus (*)
Balaenoptera musculus (*)
Tursiops truncatus (*)
Sus scrofa (.)
Manis pentadactyla (*)
Felis catus (*)
Mustela putorius
Neogale vison
Canis lupus (.)
Nyctereutes procyonoides (*)
Rhinolophus ferrumequinum
Rousettus aegyptiacus Homo
sapiens (*)
Piliocolobus tephrosceles (*)
Macaca mulatta (*)
Macaca nemestrina (*)
Choloepus didactylus
Anas platyrhynchos
Gallus gallus

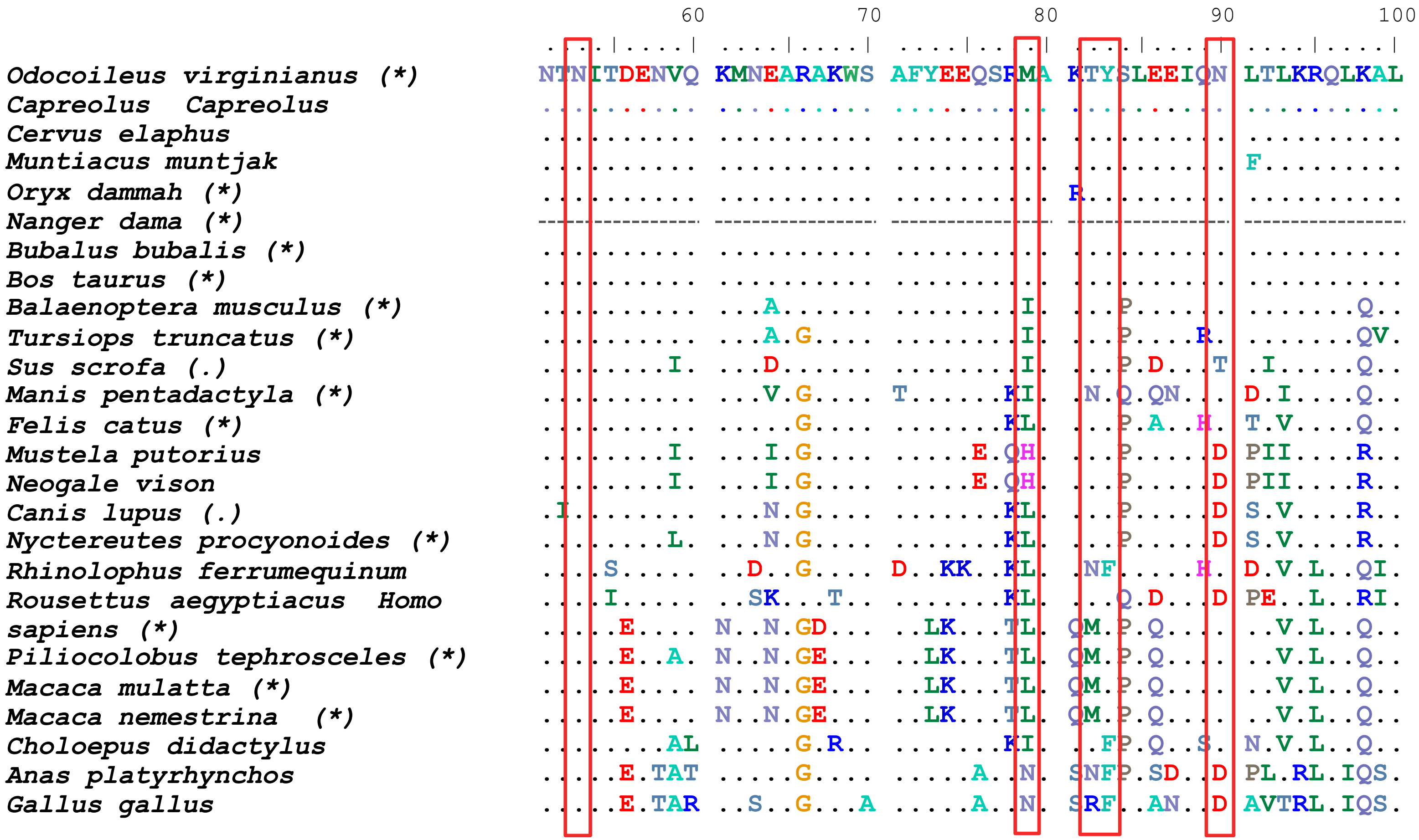
60
70
80
90
100
Odocoileus virginianus (*)
NTNITDENVQ KMNEARAKWS AFYEEQSRMA KTYSLEEIQN LTLKRQLKAL
Capreolus Capreolus
Cervus elaphus
Muntiacus muntjak
Oryx dammah (*)
Nanger dama (*)
Bubalus bubalis (*)
Bos taurus (*)
Balaenoptera musculus (*)
Tursiops truncatus (*)
Sus scrofa (.)
Manis pentadactyla (*)
Felis catus (*)
Mustela putorius
Neogale vison
Canis lupus (.)
Nyctereutes procyonoides (*)
Rhinolophus ferrumequinum
Rousettus aegyptiacus Homo sapiens (*)
Piliocolobus tephrosceles (*)
Macaca mulatta (*)
Macaca nemestrina (*)
Choloepus didactylus
Anas platyrhynchos
Gallus gallus

```
                                              110        120        130        140        150
                                      ....|....| ....|....| ....|....| ....|....| ....|....|
Odocoileus virginianus (*)            QQSGTSVLSA EKSKRLNTIL NTMSTIYSTG KVLDPN-TQE CLALEPGLDD
Capreolus  Capreolus                  ........XX XXXXX..... .......... ......-... ..........
Cervus elaphus                        .......... .......... .......... ......-... ..........
Muntiacus muntjak                     .......... .......... .......... ......-... ..........
Oryx dammah (*)                       .H.......T .......... .K........ ......-... ..........
Nanger dama (*)                       ---------- ---------- ---------- ---------- ----------
Bubalus bubalis (*)                   .H....A... ....Q..... .K........ ......-... ..........
Bos taurus (*)                        .H....A... .......... .K........ ......-... ..........
Balaenoptera musculus (*)             .......... D......... ........S. ......-... Y.V.......
Tursiops truncatus (*)                .......... D......A.. S.......S. ......-... S.V.......
Sus scrofa (.)                        ......G... D......... ........S. ......NP.. ..V......E
Manis pentadactyla (*)                .L..S.A... D.NQ...... .......... ..CN.GNP.. .SL......N
Felis catus (*)                       ....S..... D..Q...... .A........ .ACN..NP.. ..L.......
Mustela putorius                      ....S..... D.RE...... .A........ .ACN..NP.. ..L.......
Neogale vison                         ....S..... D.RE...... .A........ .ACN..NP.. ..L.......
Canis lupus (.)                       .H..S..... D.NQ...... .S...V.... .ACN.SNP.. ..L.......
Nyctereutes procyonoides (*)          .H..S..... D.NQ...... .S........ .ACN.SNP.. ..L.......
Rhinolophus ferrumequinum             ....SP...E D......S.. .A........ ..CK..NP.. ..L......N
Rousettus aegyptiacus  Homo           ....S.T... D.T....D.. .......... .ICQ..NS.. ..L.......
sapiens (*)                           ..N.S....E D......... .......... ..CN.DNP.. ..L.....NE
Piliocolobus tephrosceles (*)         ..N.S....E D......... .......... ..CN..NP.. ..L.D...NE
Macaca mulatta (*)                    ..N.S....E D......... .......... ..CN..NP.. ..L.D...NE
Macaca nemestrina  (*)                ..N.S....E D......... .......... ..CN..NP.. ..L.D...NE
Choloepus didactylus                  ....S..... D.N....I.. .A..S..... ..CNASNPK. .FF.......
Anas platyrhynchos                    .DK.S....P ..YS....V. .......... T.CKTTAPFD .MV......S
Gallus gallus                         .DR.S....P ..YS...SVM .S........ V.CKATEPFD ..V.......
```

```
                                        160        170        180        190        200
                                ....|....| ....|....| ....|....| ....|....| ....|....|
Odocoileus virginianus (*)      IMENSRDYNR RLWAWEGWRA EVGKQLRPLY EEYVVLENEM ARANNYEDYG
Capreolus  Capreolus            ........S. .......... .......... .......... ....X.....
Cervus elaphus                  .......... .......... .......... .......... ..........
Muntiacus muntjak               .......... .......... .......... .......... ..........
Oryx dammah (*)                 .......... .......... .......... .......... ..........
Nanger dama (*)                 ---------- ---------- ---------- ---------- ----------
Bubalus bubalis (*)             .......... .......... .......... .......... ..........
Bos taurus (*)                  .......... .......... .......... .......... ..........
Balaenoptera musculus (*)       .....E.... .......... ........F. .......... ..........
Tursiops truncatus (*)          .....K.... .......... .......... .......... ..........
Sus scrofa (.)                  .....K..S. ......S... .......... .......... ..........
Manis pentadactyla (*)          ...S.K...E .........S .......... ......K... ....H.....
Felis catus (*)                 .....K...E .......... .......... ....A.K... ..........
Mustela putorius                .....K...E .........S .......... ....A.K... ..........
Neogale vison                   .....K...E .........S .......... ....A.K... ..........
Canis lupus (.)                 .....K...E .........S .......... ....A.K... ..........
Nyctereutes procyonoides (*)    .....K...E .........S .......... ....A.K... ..........
Rhinolophus ferrumequinum       ...T.K...E .......... .......... ......K... ..GYH.....
Rousettus aegyptiacus  Homo     ...S.K..SQ ......S..S ........Y. ......K... ..GE......
sapiens (*)                     ..A..L...E ......S..S .......... ......K... ....H.....
Piliocolobus tephrosceles (*)   ...K.L...E .........S .......... ......K... ....H.K...
Macaca mulatta (*)              ...K.L...E .........S .......... ......K... .G..H.K...
Macaca nemestrina  (*)          ...K.L...E .........S .......... ......K... ....H.K...
Choloepus didactylus            ........DE .........S .......L.. ......K... ..........
Anas platyrhynchos              ..A..I..HE .......... D..RMM.... ....D.K..A .KL.G.A...
Gallus gallus                   ..A..I..HE .......... D..RMM.... ....E.K..A ..L...S...
```

```
                                              210        220        230        240        250
                                     ....|....| ....|....| ....|....| ....|....| ....|....|
Odocoileus virginianus (*)           DYWRGDYEVT EAGDYDYSRD QLMKDVENTF AEIKPLYEQL HAYVRAKLMD
Capreolus  Capreolus                 .......... .......... .......H.. .......... ..........
Cervus elaphus                       .......... .......... .......H.. .......... ..........
Muntiacus muntjak                    .......... .......... .......H.. .......... ..........
Oryx dammah (*)                      .......... G......... .......R.. .......... ..........
Nanger dama (*)                      ---------- ---------- ---------- ---------- ----------
Bubalus bubalis (*)                  .......... G......... .......H.. .......... .........H
Bos taurus (*)                       .......... G......... .......R.. .......... .........H
Balaenoptera musculus (*)            .......... G..V.....N ..IA...R.. .......... ..........
Tursiops truncatus (*)               .......... G......... ..IR...R.. .......... ..........
Sus scrofa (.)                       .......... GT.......N ...E...R.. ........H. ..........
Manis pentadactyla (*)               ........TE G.NG.N.... H.IE...HI. TQ......H. ..........
Felis catus (*)                      ........EE WTDG.N...S ..I....H.. TQ.....QH. ..........
Mustela putorius                     ........EE W.DG.S...N ..IE...H.. TQ......H. ..........
Neogale vison                        ........EE W.DG.N...N ..IE...H.. TQ......H. ..........
Canis lupus (.)                      ........EE WENG.N...N ..ID...L.. TQ.M...QH. .....T....
Nyctereutes procyonoides (*)         ........EE WENG.N...N ..ID...H.. TQ.M...QH. .....T....
Rhinolophus ferrumequinum            ....R...TE GSP.LE.... ..I....RI. .......... .....T....
Rousettus aegyptiacus   Homo         ........TE GINGSA.T.. ..IE..DRI. T......... .....T....
sapiens (*)                          .........N GVDG.....G ..IE...H.. E.......H. .........N
Piliocolobus tephrosceles (*)        ........AN GVDG...N.. ..IE...H.. E.......H. .........N
Macaca mulatta (*)                   .........N GVDG..NN.. ..IE...R.. E.......H. .........N
Macaca nemestrina  (*)               .........N GVDG...N.. ..IE...R.. E.......H. .........N
Choloepus didactylus                 ........TE GENG.A.N.S ..I....H.. E.......H. .....T..T.
Anas platyrhynchos                   ....AN..AD YPEE.K.... ..IQ...K.. EQ.....Q.. .....HR.EQ
Gallus gallus                        ....AN..TD YPEE.K.... ..VQ...K.. EQ.....QH. .....HR.EQ
```

```
                                        260        270        280        290        300
                                 ....|....| ....|....| ....|....| ....|....| ....|....|
Odocoileus virginianus (*)       TY-PSYISPT GCLPAHLLGD MWGRFWTNLY SLTVPFKHKP SIDVTEKMKN
Capreolus  Capreolus             ..-....... .........X .......... .......... ........E.
Cervus elaphus                   ..-....... .......... .......... .......... ........E.
Muntiacus muntjak                A.-....... .......... .......... .......... ...I....E.
Oryx dammah (*)                  ..-....... .......... .......... ......E... ..........
Nanger dama (*)                  ---------- ---------- ---------- ---------- ----------
Bubalus bubalis (*)              ..-....... .......... .......... ......E... ........E.
Bos taurus (*)                   ..-....... .......... .......... ......E... ........E.
Balaenoptera musculus (*)        A.-..R.... .......... .......... P.....GE.. .....KE.Q.
Tursiops truncatus (*)           A.-..R.... .......... .......... P.....GER. .....KE.Q.
Sus scrofa (.)                   A.-..R.... .......... .......... P.....GE.. ......A.V.
Manis pentadactyla (*)           N.-..H.... .......... .......... P.....RQ.. N....DA.V.
Felis catus (*)                  ..-..R.... .......... .......... P.....GQ.. N....DA.V.
Mustela putorius                 A.-..R.... .......... .......... P.M...RQ.. N....DA.V.
Neogale vison                    A.-..R.... .......... .......... P.M...GQ.. N....DA.V.
Canis lupus (.)                  ..-....... .......... .......... P.....GQ.. N....NA.V.
Nyctereutes procyonoides (*)     ..-....... .......... .......... P.....GQ.. N....NA.V.
Rhinolophus ferrumequinum        ..-.FH.... .......... .......... P.....GQ.. N....DA.L.
Rousettus aegyptiacus  Homo      A.-..H.... .......... ......I... P.....EQ.. N....DE.V.
sapiens (*)                      A.-......I .......... .......... ......GQ.. N....DA.VD
Piliocolobus tephrosceles (*)    A.-....... .......... .......... ......GQ.. N....DA.V.
Macaca mulatta (*)               A.-....... .......... .......... ......GQ.. N....DA.V.
Macaca nemestrina  (*)           A.-....... .......... .......... ......GQ.. N....DA.V.
Choloepus didactylus             F.-..H.... .......... .......... ......E... N....DE.VR
Anas platyrhynchos               A.GSQF..S. .......... .......... P....YPA.. N....DA.VQ
Gallus gallus                    V.GSEL.N.. .......... .......... N....YPE.. N....SA.AQ
```

| | 310 | 320 | 330 | 340 | 350 |
|---|---|---|---|---|---|
| *Odocoileus virginianus (*)* | QSWDAERIFK | EAEKFFVSIS | LPHMTQGFWD | NSMLTEPGDG | RKVVCHPTAW |
| *Capreolus Capreolus* | .......... | .......... | .......... | .......... | .......... |
| *Cervus elaphus* | .......... | .........G | .......... | .......... | .......... |
| *Muntiacus muntjak* | .........R | .......... | .......... | .......... | .......... |
| *Oryx dammah (*)* | .......... | .......... | ..Y....... | .......... | .......... |
| *Nanger dama (*)* | -......... | .......... | ..Y......E | .......... | .......... |
| *Bubalus bubalis (*)* | .......... | .......... | ..Y....... | .......... | .......... |
| *Bos taurus (*)* | .......... | .......... | ..Y....... | .......... | .......... |
| *Balaenoptera musculus (*)* | .....K.... | .........G | ..N...E..V | .......... | .......... |
| *Tursiops truncatus (*)* | .....K.... | .........G | ..N....... | .......... | .......... |
| *Sus scrofa (.)* | .....I...E | .........G | ..N......N | .......... | .......... |
| *Manis pentadactyla (*)* | .T...N.... | ........VG | ..K...T..E | .......... | .......... |
| *Felis catus (*)* | .....R.... | ........VG | ..N......E | .........S | .......... |
| *Mustela putorius* | .....R...E | ...T....VG | ..N..E...Q | .........N | .......... |
| *Neogale vison* | .....R.... | ........VG | ..N..E...Q | .........N | .......... |
| *Canis lupus (.)* | .....RK... | ........VG | ..N...E..G | .......S.S | .......... |
| *Nyctereutes procyonoides (*)* | .....RK... | ........VG | ..N......E | .......S.S | W......... |
| *Rhinolophus ferrumequinum* | .N...K.... | .........G | ..N..E...N | .....D.... | .......... |
| *Rousettus aegyptiacus* | .N.N.K.... | ........LG | ..N..ET..E | K.V....DND | Q..A...... |
| *Homo sapiens (*)* | .A...Q.... | ........VG | ..N......E | .....D..NV | Q.A....... |
| *Piliocolobus tephrosceles (*)* | .A.N.Q.... | ........VG | ..N......E | .....D...V | Q......... |
| *Macaca mulatta (*)* | .A.N.Q.... | ........VG | ..N......E | .....D..NV | Q......... |
| *Macaca nemestrina (*)* | .A.N.Q.... | ........VG | ..N......E | .....D..NV | Q......... |
| *Choloepus didactylus* | .A...K.... | ........VG | ..K....... | .........N | .N........ |
| *Anas platyrhynchos* | KN...VK... | A..A..S..G | .YN..E...K | .......T.N | .......... |
| *Gallus gallus* | KN...MK... | T..A..A..G | .YN..E...T | .......T.N | .......... |

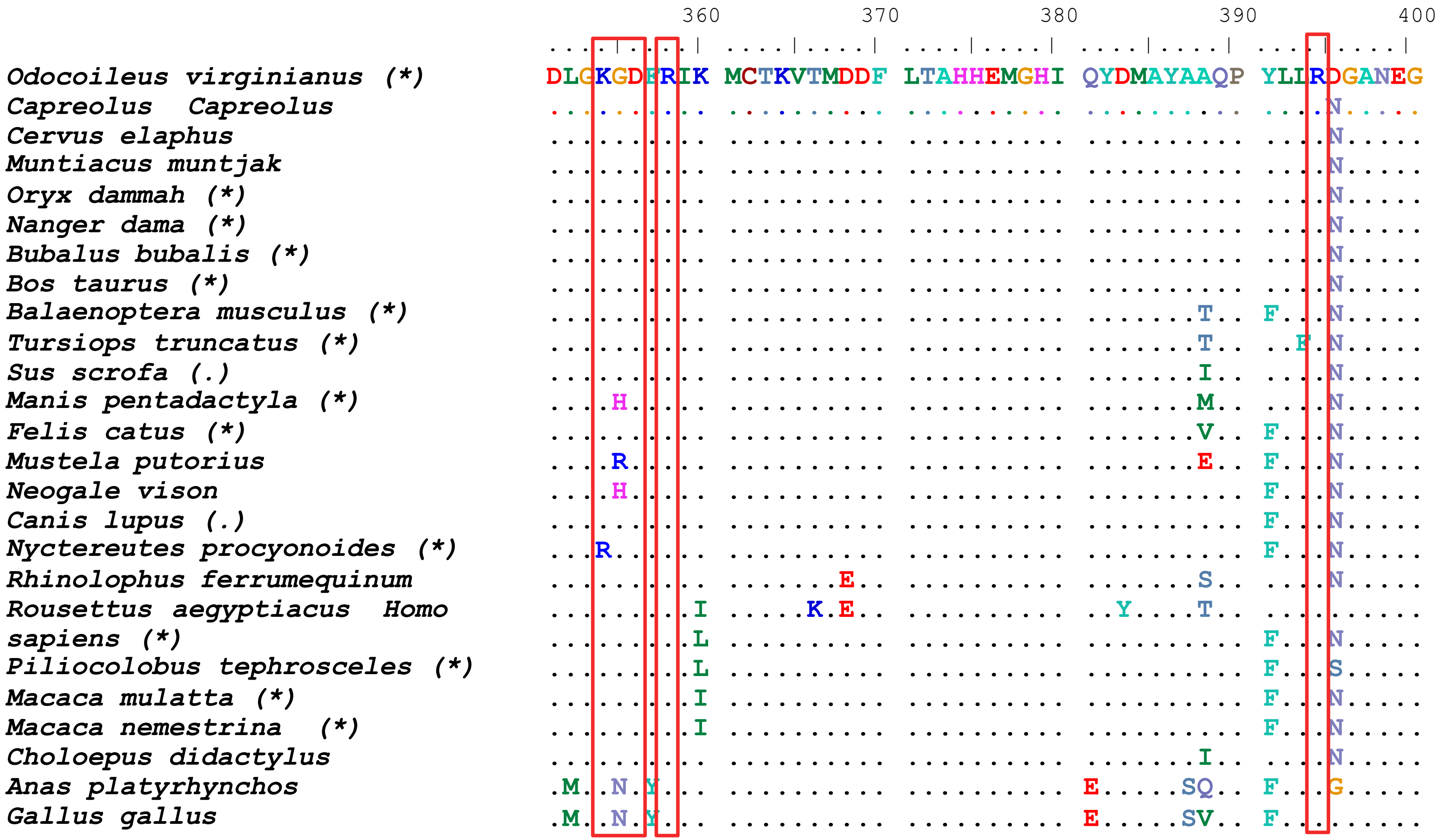

360
370
380
390
400
DLGKGDFRIK MCTKVTMDDF LTAHHEMGHI QYDMAYAAQP YLIRDGANEG
Odocoileus virginianus (*)
Capreolus Capreolus
Cervus elaphus
Muntiacus muntjak
Oryx dammah (*)
Nanger dama (*)
Bubalus bubalis (*)
Bos taurus (*)
Balaenoptera musculus (*)
Tursiops truncatus (*)
Sus scrofa (.)
Manis pentadactyla (*)
Felis catus (*)
Mustela putorius
Neogale vison
Canis lupus (.)
Nyctereutes procyonoides (*)
Rhinolophus ferrumequinum
Rousettus aegyptiacus Homo
sapiens (*)
Piliocolobus tephrosceles (*)
Macaca mulatta (*)
Macaca nemestrina (*)
Choloepus didactylus
Anas platyrhynchos
Gallus gallus

```
                                              410        420        430        440        450
                                    ....|....| ....|....| ....|....| ....|....| ....|....|
Odocoileus virginianus (*)          FHEAVGEIMS LSAATPHYLK ALGLLEPDFY EDNETEINFL LKQALTIVGT
Capreolus  Capreolus                .......... .......... .......... .......... ..........
Cervus elaphus                      .......... .......... .......... .......... ..........
Muntiacus muntjak                   .......... .......... .......... .......... ..........
Oryx dammah (*)                     .......... .......... .....A.... .......... ..........
Nanger dama (*)                     .......... .......... .....A.... .......... ..........
Bubalus bubalis (*)                 .......... .......... .....A...H .......... ..........
Bos taurus (*)                      .......... .......... .....A...H .......... ..........
Balaenoptera musculus (*)           .......... .......... .....P.... ...V...... .....Q....
Tursiops truncatus (*)              .......... .......... .....P.... ..SA...... ..........
Sus scrofa (.)                      .......... .......... .....P.... ..S....... ..........
Manis pentadactyla (*)              .......... ......KH.. NI...P.... .......... ..........
Felis catus (*)                     .......... ......NH.. TI...S.G.S ..S....... ..........
Mustela putorius                    .......... ......NH.. NI...P...S ..S..D.... ..........
Neogale vison                       .......... ......NH.. NI...P...S ..S..D.... ..........
Canis lupus (.)                     .......... ......NH.. NI...P.S.F ..S....... ..........
Nyctereutes procyonoides (*)        .......... ......NH.. NI...P.S.F ..S....... ..........
Rhinolophus ferrumequinum           .......V.. ..V...KH.. TM...SS..L .......... F....N....
Rousettus aegyptiacus  Homo         .......VI. ..V...NH.. NM...P.... .......... .....NV...
sapiens (*)                         .......... ......KH.. SI...S...Q .......... ..........
Piliocolobus tephrosceles (*)       .......... ......KH.. SI...S...Q .......... ..........
Macaca mulatta (*)                  .......... ......KH.. SI...S...Q .......... ..........
Macaca nemestrina  (*)              .......... ......KH.. SI...S...Q .......... ..........
Choloepus didactylus                .......... ......KH.. .I...P...Q ..F....... ..........
Anas platyrhynchos                  .......... ......EH.. S.D....T.Q ..E....... ..........
Gallus gallus                       .......... ......QH.. S.D....T.Q ..E....... ..........
```

```
                                          460        470        480        490        500
                                 ....|....| ....|....| ....|....| ....|....| ....|....|
Odocoileus virginianus (*)       LPFTYMLEKW RWMVFKGEIP KEQWMEKWWE MKREIVGVVE PLPHDETYCD
Capreolus  Capreolus             .......... .......... .......... .......... ..........
Cervus elaphus                   .......... .......... .....Q.... .......... ..........
Muntiacus muntjak                .......... .......... .....Q.... .......... ..........
Oryx dammah (*)                  .......... .......... .Q........ .......... ..........
Nanger dama (*)                  .......... .......... .Q........ .......... ..........
Bubalus bubalis (*)              .......... .......... .Q........ .......... ..........
Bos taurus (*)                   .......... .......... .Q........ .......... ..........
Balaenoptera musculus (*)        .......... .......... .....Q.... .......... ..........
Tursiops truncatus (*)           .......... .......... .....Q.... .......... ..........
Sus scrofa (.)                   .......... .......... .....Q.... .......... ..........
Manis pentadactyla (*)           .......... .....S.Q.. .....K.... .......... .V........
Felis catus (*)                  .......... .......... .....Q.... .......... .V........
Mustela putorius                 .......... .......... .....Q.... ...D...... ..........
Neogale vison                    .......... .......... .....Q.... ...D...... ..........
Canis lupus (.)                  .......... .......... .D...KT... ...N...... .V........
Nyctereutes procyonoides (*)     .......... .......... .D...KT... ...N...... .V........
Rhinolophus ferrumequinum        .......... .......... ..E..K.... ...K...... .V........
Rousettus aegyptiacus  Homo      .......... .......... .......... ....L..... ..........
sapiens (*)                      .......... .......... .D...K.... .......... .V........
Piliocolobus tephrosceles (*)    .......... .....E.... .D...K.... .......... .V........
Macaca mulatta (*)               .......... .......... .D...K.... .......... .V........
Macaca nemestrina  (*)           .......... .......... .D...K.... .......... .V........
Choloepus didactylus             .......... .....R.... ....TK.... ..Q....M.. .V...S....
Anas platyrhynchos               M......... .....R...T .QE.TKQ... ...D...... .V........
Gallus gallus                    M......... .....N...T .QE.TKR..K .......... .V........
```

```
                                                  510        520        530        540        550
                                         ....|....|  ....|....|  ....|....|  ....|....|  ....|....|
Odocoileus virginianus (*)               PACLFHVAED  YSFIRYYTRT  IYQFQFHEAL  CKTANHEGAL  FKCDISNSTE
Capreolus  Capreolus                     ..........  ..........  ..........  ..........  ..........
Cervus elaphus                           ..........  ..........  ..........  ....K.....  ..........
Muntiacus muntjak                        ..........  ..........  ..........  ..........  ..........
Oryx dammah (*)                          ..........  ..........  ..........  ....K.....  ..........
Nanger dama (*)                          ..........  ..........  ..........  ....K.....  ..........
Bubalus bubalis (*)                      ..........  ..........  ..........  ....K.....  ..........
Bos taurus (*)                           ..........  ..........  ..........  ....K.....  ..........
Balaenoptera musculus (*)                ..........  ..........  ..........  .Q..K...P.  Y.........
Tursiops truncatus (*)                   ..........  ..........  ..........  .Q..K...P.  Y.........
Sus scrofa (.)                           ..........  ..........  ..........  .R..K...P.  Y.........
Manis pentadactyla (*)                   ..S.....N.  ..........  ......Q...  .Q..K...P.  H.........
Felis catus (*)                          ..S.....N.  ..........  ......Q...  .RI.K...P.  H.......S.
Mustela putorius                         ..A.....N.  ..........  ......Q...  .QI.K...P.  Y.......S.
Neogale vison                            ..A.....N.  ..........  ......Q...  .QI.K...P.  Y.......R.
Canis lupus (.)                          ..S.....N.  ..........  ......Q...  .QI.K...P.  H.......S.
Nyctereutes procyonoides (*)             ..S.....N.  ..........  ......Q...  .QI.K...P.  H.......S.
Rhinolophus ferrumequinum                ..S.....N.  ..........  .FE.......  .RI.K.D.P.  H........D
Rousettus aegyptiacus  Homo              ..S.....N.  ..........  .FE...L...  .RI.Q...P.  Y...A.....
sapiens (*)                              ..S....SN.  ..........  L.....Q...  .QA.K...P.  H.........
Piliocolobus tephrosceles (*)            ..S....SN.  ..........  L.....Q...  .QA.K...P.  H.........
Macaca mulatta (*)                       ..S....SN.  ..........  L.....Q...  .QA.K...P.  H.........
Macaca nemestrina  (*)                   ..S....SN.  ..........  L.....Q...  .QA.K...P.  H.........
Choloepus didactylus                     ..T.....N.  ..........  ......Q...  .QA...Q.P.  HR........
Anas platyrhynchos                       ..A.....N.  ..........  ..........  ..A...T.P.  HT...T...A
Gallus gallus                            ..A.....N.  ..........  ......Q...  ..A...T.P.  H....T...A
```

```
                                       560        570        580        590        600
                               ....|....| ....|....| ....|....| ....|....| ....|....|
Odocoileus virginianus (*)     AGQRLLQMLS LGKSEPWTLA LESIVGIKTM DVKPLLNYFE PLFTWLKEQN
Capreolus  Capreolus           .......... .......... .......... .......... ..........
Cervus elaphus                 .......... .......... .......... .......... ..........
Muntiacus muntjak              .......... .......... .......... .......... ..........
Oryx dammah  (*)               .........R .......... .......... .......... ..........
Nanger dama  (*)               .........R .......... ..N....... .......... ..........
Bubalus bubalis  (*)           .........R .......... ..N....... .......... ..........
Bos taurus  (*)                .........R .......... ..N....... .......... ..........
Balaenoptera musculus  (*)     .........H .......... ..N...V... .......... ..L.......
Tursiops truncatus  (*)        .........H ........S. ..R...V... .......... ..L....G..
Sus scrofa  (.)                ...K...... .......... ..N...V... ......S... ..L....A..
Manis pentadactyla  (*)        ...K...... ....K..... ..RV..T.N. ..R....... ..L.......
Felis catus  (*)               ..KK.....T ....K..... ..HV..E.K. N.T...K... ..........
Mustela putorius               ...K.HE... ..R.K...F. ..RV..A... ..R....... ..........
Neogale vison                  ...K.HE... ..R.K...F. ..RV..A... ..R....... ..........
Canis lupus  (.)               ...K..E..K ....K...Y. ..IV..A.N. ..R....... ..........
Nyctereutes procyonoides  (*)  ...K..E..K ....K...Y. ..IV..A.N. ..R....... ..........
Rhinolophus ferrumequinum      ..EK.H.... V...Q...SV .KDF..S.N. ..G...R... ..Y...T...
Rousettus aegyptiacus  Homo    ..KK.H.... ....K..... ....A.T.N. ..R....... ........K.
sapiens (*)                    ...K.FN..R .......... ..NV..A.N. N.R....... .......D..
Piliocolobus tephrosceles (*)  ...K..N..K .......... ..NV..A.N. N.R....... .......D..
Macaca mulatta  (*)            ...K..N..K ..E....... ..NV..A.N. N.R....... .......D..
Macaca nemestrina   (*)        ...K..N..K .......... ..NV..A.N. N.R....... .......D..
Choloepus didactylus           ...K..N..K S.......A. ..HV..T.H. .......... .......D..
Anas platyrhynchos             ..GS.REL.K ..R.K...Q. ...LT.E.Y. NAT...H... ...N..QKN.
Gallus gallus                  ..GN.R.L.E ....K...Q. ...AT.E.Y. NAT...H... ...N..QKN.
```

```
                                        610        620        630        640        650
                                ....|....| ....|....| ....|....| ....|....| ....|....|
Odocoileus virginianus (*)      RNSFVGWSTE WTPYSDQSIK VRIS------ ---------- ---LKSALG-
Capreolus  Capreolus            .......... ...XXXXXXX XXXXXXXXXX XXXYEWNDNE MYLFR..VAY
Cervus elaphus                  K......... .......... ....LKSALG KNAYEWNDNE LYLFR.SVAY
Muntiacus muntjak               .......... .......... ....LKSALG KNAYEWNDNE MYLFR.SVAY
Oryx dammah (*)                 .......... .......... ....LKSGLG KNAYEWNDNE MYLFR.SVAY
Nanger dama (*)                 .......... .......... ....LKSALG KNAYEWNDNE MYLFR.SVAY
Bubalus bubalis (*)             .......... .......... ....LKAALG ENAYEWNDNE MYLFR.SVAY
Bos taurus (*)                  .......... .......... ....LKSALG ENAYEWNDNE MYLFQ.SVAY
Balaenoptera musculus (*)       ...S.....D .......... ....LKSALG EKAYEWNDNE MYLFR.SVAY
Tursiops truncatus (*)          .......R.D .....N.... ....LKSALG EKAYEWNDNE MYLFR.SVAY
Sus scrofa (.)                  G..S...N.D ....A..... ....LKSALG KEAYEWNDNE MYLFR.SIAY
Manis pentadactyla (*)          K......N.D .S..AA.... ....LKSALG EKAYEWNDSE MYLFR.SVAY
Felis catus (*)                 .......N.D .R..A..... ....LKSALG DEAYEWNDNE MYLFR.SVAY
Mustela putorius                .......N.D .S..A..... ....LKSALG EKAYEWNDNE MYFFQ.SIAY
Neogale vison                   .......N.D .S..A..... ....LKSALG EKAYEWNDNE MYFFQ.SIAY
Canis lupus (.)                 .......N.D .S..A..... ....LKSALG EKAYEWNNNE MYLFR.SIAY
Nyctereutes procyonoides (*)    .......N.D .S..A..... ....LKSALG EKAYEWNNNE MYLFR.SIAY
Rhinolophus ferrumequinum       .K.....N.D .S..A..... ....LKSALG EKAYEWNNNE MYLFR.SVAY
Rousettus aegyptiacus  Homo     .........D .S...G.... ....LKAALG EKAYEWNDNE MYLF..SIAY
sapiens (*)                     K........D .S..A..... ....LKSALG DKAYEWNDNE MYLFR.SVAY
Piliocolobus tephrosceles (*)   K........D .S..A..... ....LKSALG DKAYEWNDNE MYLFR.SVAY
Macaca mulatta (*)              K........D .S..A..... ....LKSALG DKAYEWNDNE MYLFR.SVAY
Macaca nemestrina  (*)          K........D .S..A..... ....LKSALG DKAYEWNDNE MYLFR.SVAY
Choloepus didactylus            ..VP.....D GS.DA..-.. ....LKSALG DKAYEWNDNE MYLFR.SVAY
Anas platyrhynchos              SGRYI..N.D .....ENA.. ....LKAAG- -QTYEWNKSE LFLF..TIAY
Gallus gallus                   SGRSI..N.D ......NA.. ....LKAALG DDAYVWDASE LFLF..SIAY
```

```
                                     660        670        680        690        700
                               ....|....| ....|....| ....|....| ....|....| ....|....|
Odocoileus virginianus (*)     ---------- ---------- ---KNADANC PFVWCVPPVS HLVAIVIRSA
Capreolus  Capreolus           AMRKYFLGER NETIPFGEEN VWVSDKKPRI S.KFF.TS.N NVSD..P.TE
Cervus elaphus                 AMRKYFLKKR NETIPFGEEN VWVSDKKPRI S.KFF.TSPN NVSD.IP.TE
Muntiacus muntjak              AMRKYFLKKR NETIPFGEEN VWVSDKKPRI S.KFF.TSPN NVSD.IP.TE
Oryx dammah (*)                AMRKYFFKDR NETIPFGEEN VWVSDKKPRI S.KFF.TSPN NVSD.IP.TE
Nanger dama (*)                AMRRYFFEAS NETIPFGAEN VWVSDKKPRI S.KFF.TSPN NVSD.IP.TE
Bubalus bubalis (*)            AMRKYFSEAR NETVLFGEDN VWVSDKKPRI S.KFF.TSPN NVSD.IP.TE
Bos taurus (*)                 AMRKYFSEAR NETVLFGEDN VWVSDKKPRI S.KFF.TSPN NVSD.IP.TE
Balaenoptera musculus (*)      AMREYFSKVR NETIPFGEKD VWVSDLKPRI S.NFF.TTPK NVSD.IS.TE
Tursiops truncatus (*)         AMREYFSKVR NKTIPFGEKD VWVSDLKPRI S.NFF.TSPK NMSD.IP.TE
Sus scrofa (.)                 AMRNYFSSAK NETIPFGAED VWVSDLKPRI S.NFF.TSPA NMSD.IP..D
Manis pentadactyla (*)         AMREYFSKFK KQTIPFEEES VRVSDLKPRV S.IFF.TLPK NVS.VIP.AE
Felis catus (*)                AMREYFSKVK NQTIPFVEDN VWVS.LKPRI S.NFF.TASK NVSDVIP..E
Mustela putorius               AMREYFSKVK NQTIPFVGKD VRVSDLKPRI S.NFI.TSPE NMSD.IP.AD
Neogale vison                  AMREYFSKVK KQTIPFVDKD VRVSDLKPRI S.NFI.TSPE NMSD.IP.AD
Canis lupus (.)                AMRQYFSEVK NQTIPFVEDN VWVSDLKPRI S.NFS.TSPG NVSD.IP.TE
Nyctereutes procyonoides (*)   AMRQYFSEVK NQTIPFVEDN VWVSDLKPRI S.NFF.TSPG NVSD.IP.TE
Rhinolophus ferrumequinum      AMREYFLKTK NQTILFGEED VWVS.LKPRI S.NFY.TSPR N.SD.IP.PE
Rousettus aegyptiacus  Homo    SLREYFLKVK NLTIPFGEED VWVSDLKPRI S.NFF.TSPQ NVSEFIP.TE
sapiens (*)                    AMRQYFLKVK NQMILFGEED VRVA.LKPRI S.NFF.TAPK NVSD.IP.TE
Piliocolobus tephrosceles (*)  AMRKYFLEIK HQTILFGEED VRVADLKPRI S.NFY.TAPK NVSD.IP.TE
Macaca mulatta (*)             AMRTYFLEIK HQTILFGEED VRVADLKPRI S.NFY.TAPK NVSD.IP.TE
Macaca nemestrina  (*)         AMRTYFLEIK HQTILFGEED VRVADLKPRI S.NFY.TAPK NVSD.IP.TE
Choloepus didactylus           AMREYFLKVK NQTILFGEED VQVSELKPRI S.IFV.SAPN NTSD.IP.TE
Anas platyrhynchos             AMRTYFAQ-K QQLIDFEATD VHVSEETQRV S.YIT.SMPG NASN..PKAD
Gallus gallus                  AMRKYFAKEK EQNVDFQVTD IHVGEETQRV S.YLT.SMPG NVSD..P.AD
```

```
                                           710        720        730        740        750
                                   ....|....| ....|....| ....|....| ....|....| ....|....|
Odocoileus virginianus (*)         VT_________ __________ __________ __________ _________V
Capreolus  Capreolus               .EEAIRLSRG RINDAFQLDD NSLEFLGIQP TLGPPYEPPV TIWLIIFGV.
Cervus elaphus                     .ENAIRLSRY RINDAFQLDD NSLEFLGIQP TLGPPYKPPV TIWLIIFGV.
Muntiacus muntjak                  .ENAIRLSRD RINDAFQLDD NSLEFLGIQP TLGPPYEPPV TIWLIIFGV.
Oryx dammah (*)                    .ENAIRLSRD RINDAFQLDD NSLEFLGIQP TLGPPYEPPV TIWLIIFGV.
Nanger dama (*)                    .ENAIRLSRD RINDAFQLDD NSLEFLGIQP TLGPPYEPPV TIWLIIFGV.
Bubalus bubalis (*)                .ENAIRLFRG RINDVFQLDD NSLEFLGIQP TLRPPYEPPV TIWLIIFGV.
Bos taurus (*)                     .ENAIRLSRD RFNDVFQLDD NSLEFLGIQP TLGPPYEPPV TIWLIIFGV.
Balaenoptera musculus (*)          .EEAIRMSRG RINDAFRLDD NSLEFLGIQP TLGPPYEPPV TIWLIIFGA.
Tursiops truncatus (*)             .EEAIRMSRG RINDAFRLDD SSLEFLGVQP TLAPPYEPPV TVWLIIFGV.
Sus scrofa (.)                     .EKAISMSRS RINDAFRLDD NTLEFLGIQP TLGPPDEPPV TVWLIIFGV.
Manis pentadactyla (*)             .EEAIRMSRS RINDVFRLDD NSLEFLGIQP TLEPPYQPPV TIWLIVFGV.
Felis catus (*)                    .EEAIRMSRS RINDAFRLDD NSLEFLGIQP TLSPPYQPPV TIWLIVFGV.
Mustela putorius                   .EEAIRKSRG RINDAFRLDD NSLEFLGIQP TLEPPYQPPV TIWLIVFGV.
Neogale vison                      .EEAIRKSRG RINDAFRLDD NSLEFLGIQP TLEPPYQPPV TIWLIVFGV.
Canis lupus (.)                    .EEAIRMYRS RINDVFRLDD NSLEFLGIQP TPGPPYEPPV TIWLIVFGV.
Nyctereutes procyonoides (*)       .EEAIRMYRS RINDVFRLDD NSLEFLGIQP TLGPPYEPPV TIWLIVFGV.
Rhinolophus ferrumequinum          .EGAIRMSRS RINDAFRLDD NSLEFLGIQP TLGPPYQPPV TIWLIVFGV.
Rousettus aegyptiacus  Homo        .EGAIRMSRS RINDAFRLDD DTLEFLGIEP TLGTPYQPPV TIWLIVFGV.
sapiens (*)                        .EKAIRMSRS RINDAFRLND NSLEFLGIQP TLGPPNQPPV SIWLIVFGV.
Piliocolobus tephrosceles (*)      .EEAIRLSRS RINDAFRLND DSLEFLGIQP TLAPPYQPPV TIWLIVFGV.
Macaca mulatta (*)                 .EEAIRISRS RINDAFRLND NSLEFLGIQT TLAPPYQSPV TTWLIVFGV.
Macaca nemestrina  (*)             .EEAIRISRS RINDAFRLND NSLEFLGIQT TLAPPYQSPV TTWLIVFGV.
Choloepus didactylus               .EKAISMSRG RINDAFRLDD NTLEFVGIYP TLAPPYEPPV VIWLVVFGVI
Anas platyrhynchos                 .ESAISMSRG RINEAFGLDD DTLEFVGIIP TLAAPYEPPV TIWLIIFGV.
Gallus gallus                      .EKAIRMSRG RISEAFRLDD NTLEFDGIVP TLATPYKPPV TIWLILFGV.
```

```
                                          760        770        780        790        800
                                 ....|....| ....|....| ....|....| ....|....| ....|....|
Odocoileus virginianus (*)       SQCCVQATLV LLNPGPK--- -----VPEE- ---------- ----------
Capreolus  Capreolus             MGVV.LGIV. .IFT.IRDRR KKNQASS..N PYG------- SVDLNK--GE
Cervus elaphus                   MGVV.IGIV. .IFT.IRDRR KKNQASS..N PYG------- SVDLNK--GE
Muntiacus muntjak                MGVV.IGIV. .IFT.IRDRR KKNQVSS..N PYG------- SVDLNK--GE
Oryx dammah (*)                  MGVV.IGIVG .IFT.IRDQR KKNQASS..N PYG------- SVDLNK--GE
Nanger dama (*)                  MGVV.IGIV. .IFT.IQDRR KHDNDGLQND ENLRVQQQAV KVDIPR--NS
Bubalus bubalis (*)              MGVV.IGII. .IFT.IRDRR KKNQASS..N PYG------- SVDLNK--GE
Bos taurus (*)                   MGVV.IGIV. .IFT.IRNRR KKNQASS..N PYG------- SVDLNK--GE
Balaenoptera musculus (*)        MGVV.IGIA. .IFT.IRDRR EKSQASS..N PYI------- SMDLSK--GE
Tursiops truncatus (*)           MGVV.IGIV. .IFT.IRDRR KKNQASS..N PYG------- SVGLSK--GE
Sus scrofa (.)                   MGLV.VGIV. .IFT.IRDRR KKKQASS..N PYG------- SMDLSK--GE
Manis pentadactyla (*)           MGVI.VGIV. .IFT.IRDRK KKNQARS.QN PYA------- SVDLSK--GE
Felis catus (*)                  MGVV.VGIVL .IVS.IRNRR KNNQARS..N PYA------- SVDLSK--GE
Mustela putorius                 MGVV.VGIFL .IFS.IRNRR KNNQARS..N PYA------- SVDLSK--GE
Neogale vison                    MGVV.VGIFL .IFS.IRNRR KNNQARS..N PYA------- SVDLSK--GE
Canis lupus (.)                  MGVV.VGIVL .IFS.IRNRR KNDQARG..N PYA------- SVDLSK--GE
Nyctereutes procyonoides (*)     MGVV.VGIVL .IFS.IRNRR KNDQARG..N PYA------- SVDLSK--GE
Rhinolophus ferrumequinum        MAVV.VGIV. .IIT.IRDRR KKDQARS..N PYS------- SVDLSK--GE
Rousettus aegyptiacus  Homo      MGLV.VGIVL .IFV.IRDRR KKNQERS..N PYS------- SVDLSK--GE
sapiens (*)                      MGVI.VGIVI .IFT.IRDRK KKNKARSG.N PYA------- SIDISK--GE
Piliocolobus tephrosceles (*)    MGVI.AG.V. .IFT.IRDRK KKNQARS..N PYA------- SIDISK--GE
Macaca mulatta (*)               MGVI.AGIV. .IFT.IRDRK KKNQARS..N PYA------- SIDINK--GE
Macaca nemestrina  (*)           MGVI.AGIV. .IFT.IRDRK KKNQARS..N PYA------- SIDINK--GE
Choloepus didactylus             MSVI.IGIVL .IFT.IRERK KNRQASGQ.N PYA------- SVDLSK--GE
Anas platyrhynchos               ISLV.IGVI. .IVS.QRDRK KKAKGRER.A ESN------C EVNPYDDDGR
Gallus gallus                    MSLI.IGVI. .IIT.QRDKR KKARGRAN.A GSN------C EVNPYDEDGR
```

| | |
|---|---|
| | 810 |
| | ....\|....\| ....\| |
| *Odocoileus virginianus (*)* | ---------- ----- |
| *Capreolus Capreolus* | NNSGFQNTDD VQTSL |
| *Cervus elaphus* | NNSGFQNTDD VQTSL |
| *Muntiacus muntjak* | NNSGFQNTDD VQTSL |
| *Oryx dammah (*)* | NNSGFQNTDD VQTSL |
| *Nanger dama (*)* | LKATVPFSNS PEKLK |
| *Bubalus bubalis (*)* | NNSGFQNTDD VQTSL |
| *Bos taurus (*)* | NNSGFQNIDD VQTSL |
| *Balaenoptera musculus (*)* | NNSGFQNSGD VHTSF |
| *Tursiops truncatus (*)* | NNPGFQNSDD VQTSF |
| *Sus scrofa (.)* | SNSGFQNGDD IQTSF |
| *Manis pentadactyla (*)* | NNPGFQNVDD VQTSF |
| *Felis catus (*)* | NNPGFQHADD VQTSF |
| *Mustela putorius* | NNPGFQNVDD VQTSF |
| *Neogale vison* | NNPGFQNVDD VQTSF |
| *Canis lupus (.)* | NNPGFQSGDD VQTSF |
| *Nyctereutes procyonoides (*)* | NNPGFQNVDD AQTSF |
| *Rhinolophus ferrumequinum* | NNPGFQNGDD VQTSF |
| *Rousettus aegyptiacus Homo* | NNAGFQNNDD VQTSF |
| *sapiens (*)* | NNPGFQNTDD VQTSF |
| *Piliocolobus tephrosceles (*)* | NNPGFQNTDD VQTSF |
| *Macaca mulatta (*)* | NNPGFQNTDD VQTSF |
| *Macaca nemestrina (*)* | NNPGFQNTDD VQTSF |
| *Choloepus didactylus* | NNPGFQNSDD IQTSF |
| *Anas platyrhynchos* | SNKGFELSDE TQTSF |
| *Gallus gallus* | SNKGFEQSEE TQTSF |
